# mbctools: A User-Friendly Metabarcoding and Cross-Platform Pipeline for Analyzing Multiple Amplicon Sequencing Data across a Large Diversity of Organisms

**DOI:** 10.1101/2024.02.08.579441

**Authors:** Christian Barnabé, Guilhem Sempéré, Vincent Manzanilla, Joel Moo Millan, Antoine Amblard-Rambert, Etienne Waleckx

**Author notes:** Co–first authors, contributed equally (Christian Barnabé, Guilhem Sempéré).

## Abstract

We developed a python package called *mbctools*, designed to offer a cross-platform tool for processing amplicon data from various organisms in the context of metabarcoding studies. It can handle the most common tasks in metabarcoding pipelines such as paired-end merging, primer trimming, quality filtering, sequence denoising, zero-radius operational taxonomic unit (ZOTU) filtering, and has the capability to process multiple genetic markers simultaneously. *mbctools* is a menu-driven program that eliminates the need for expertise in command-line skills and ensures documentation of each analysis for reproducibility purposes. The software, designed to run in a console, offers an interactive experience, guided by keyboard inputs, assisting users along the way through data processing and hiding the complexity of command lines by letting them concentrate on selecting parameters to apply in each step of the process. In our workflow, VSEARCH is utilized for processing *fastq* files derived from amplicon-based Next-Generation Sequencing data. This software is a versatile open-source tool for processing amplicon sequences, offering advantages such as high speed, efficient memory usage, and the ability to handle large datasets. It provides functions for various tasks such as dereplication, clustering, chimera detection, and taxonomic assignment. VSEARCH is thus very efficient in retrieving the overall diversity of a sample. To adapt to the diversity of projects in metabarcoding, we facilitate the reprocessing of datasets with the possibility to adjust parameters. *mbctools* can also be launched in a headless mode, making it suited for integration into pipelines running on High-Performance Computing environments. *mbctools* is available at https://github.com/GuilhemSempere/mbctools, https://pypi.org/project/mbctools/.

## Introduction

Metabarcoding consists in identifying a targeted subset of genomes within bulk samples by massive sequencing of amplicons of taxonomically informative genetic markers, also known as barcodes (Figure 1) (Taberlet et al., 2012). Over the past decade, this approach has quickly gained popularity since Hebert et al. (2003) first advocated the use of short variable DNA sequences, amplified using universal primers, for species identification, and discovery of new taxa. Its simplicity of implementation enables biologists, local governments, and NGOs to increase their understanding of spatial and temporal ecological networks (Holdaway et al., 2017). Using universal primers specific of kingdoms or subgroups of organisms of interest (e.g. “arthropods”, “vertebrates”, “reptiles”, “amphibians”, “mammals”, “birds”, …), metabarcoding is highly effective for species-level identification and assessment of the whole diversity of a sample within the targeted kingdoms or subgroups of organisms of interest. For example, universal primers for vertebrates, targeting the 12S rRNA gene, have been proposed for metabarcoding studies focused on vertebrates (Riaz et al., 2011). In studies targeting animals, primers allowing the amplification of a portion of the mitochondrial marker Cytochrome oxidase 1 (COI) may be used (Hebert et al., 2016). In plants, standard DNA barcoding generally involves one to four plastid DNA regions (rbcL, matK, trnH-psbA, trnL), sometimes in combination with the internal transcribed spacers of nuclear ribosomal DNA (nrDNA, ITS) (CBOL Plant Working Group et al., 2009). For fungi, the internal transcribed spacer (ITS) regions ITS1 and ITS2 of the nuclear ribosomal DNA are the most commonly used barcodes (Schoch et al., 2012). For bacterial and archaeal communities, the 16S ribosomal RNA (16S rRNA) is the most widely used barcode and provides taxonomic resolution to the genus level (Caporaso et al., 2011). Beyond these markers, specific markers are used in various fields to genotype species. For instance, in the study of *Trypanosoma cruzi*, the causative agent of Chagas disease, the Glucose-6-Phosphate Isomerase (GPI) and the Cytochrome oxidase 2 (COII) genes are frequently used to distinguish between different strains (Lauthier et al., 2012; López-Dominguez et al., 2022; Barnabé et al., 2023). Although single markers are very informative, in many cases, a combination of markers is necessary to fully understand the spatial and temporal dynamics of ecological networks. A huge advantage of metabarcoding is that it is based on massive sequencing. As a consequence, all the amplicons amplified from different targeted markers can be pooled and sequenced simultaneously (Hernández-Andrade et al., 2019). This powerful approach has proven to be useful for diet profile analysis, air and water quality monitoring, biodiversity monitoring, and food quality control, e.g., beverage or ancient ecosystem composition (Leray & Knowlton, 2015; Thomsen & Willerslev, 2015; Raclariu et al., 2017). As another application example, our group has recently proposed to use this approach to untangle the transmission cycles of vector-borne pathogens and their dynamics, using the gut contents of vector insects as bulk samples, and simultaneously sequencing markers allowing for the molecular identification of vector species and/or genetic diversity, blood meal sources (using universal primers for vertebrates which may also serve as pathogen hosts), gut microbiome composition (which may modulate vectorial capacity), and pathogen diversity. Indeed, all the latter components interact together to shape transmission cycles (Dumonteil et al., 2018; Hernández-Andrade et al., 2019).

**Figure 1.**
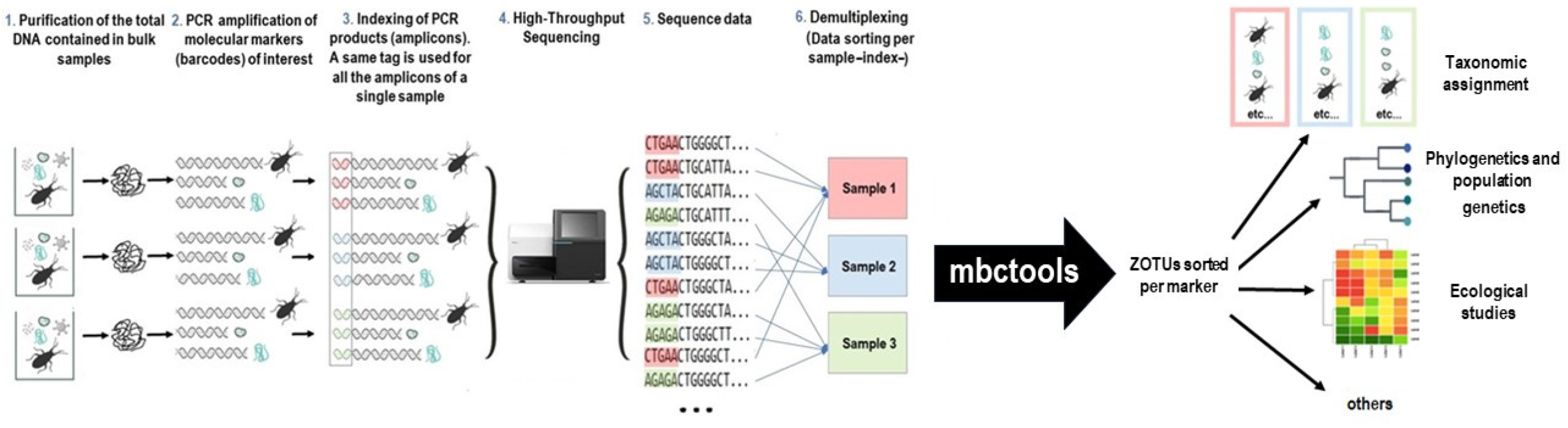
Metabarcoding workflow example and area of operation of mbctools

The increasing demand for efficient and versatile pipelines to analyze amplicon data has led to the development of over 30 programs and pipelines in recent years (Hakimzadeh et al., 2023). However, these applications often present some challenges in terms of reproducibility and user-friendliness for non-expert users in the field of bioinformatics. Indeed, the obtaining of pseudo quantitative taxonomic data from fastq files involves a series of complex processing steps that can be achieved with specific software implying the understanding of a large panel of parameters. To address this problem, mbctools was designed with a simplified interface, allowing scientists without advanced scripting skills to easily install, navigate, utilize the pipeline while focusing on key parameters. This accessibility ensures that non-expert users can effectively and autonomously analyze their data without experiencing difficulties in reproducibility or requiring extensive bioinformatics knowledge.

Among pipelines and algorithms used in the field of metabarcoding data processing, substantial variations in sensitivity and specificity are observed (Bailet et al., 2020; Mathon et al., 2021). From these benchmarking studies, VSEARCH appears appropriate for our applications. mbctools’ main purpose is to facilitate the use of VSEARCH, which has demonstrated strong performance in handling large-scale next-generation sequencing (NGS) data (Rognes et al., 2016), and it thus could be considered a “wrapper”, hiding the complexity of command-line to the users. It thus allows them to easily process sequencing reads and obtain meaningful outputs, i.e, zero-radius operational taxonomic units (ZOTUs (Edgar, 2016), almost equivalent to ASVs (amplicon sequence variants)) along with abundance information.

*mbctools* processes demultiplexed metabarcoding data obtained from one or multiple genetic markers to distribute cleaned and filtered sequences into ZOTUs sorted per marker, which can be used for taxonomic assignment or further analyzes such as phylogeny (Figures 1, 2). While the original goal in developing *mbctools* was providing assistance to novice users through its interface, it can also prove useful to bioinformaticians willing to integrate it as part of a wider data processing pipeline, since its series of mandatory operations (initial analysis described below) can be launched in a headless mode by feeding the program with a configuration file. Additionally, addressing the challenge of processing large numbers of markers simultaneously across different kingdoms was one of the main motivations for designing *mbctools*, which allows for the simultaneous processing of multiple genetic markers and has a simplified graphical interface that guides the user through key parameters of the analysis.

**Figure 2.**
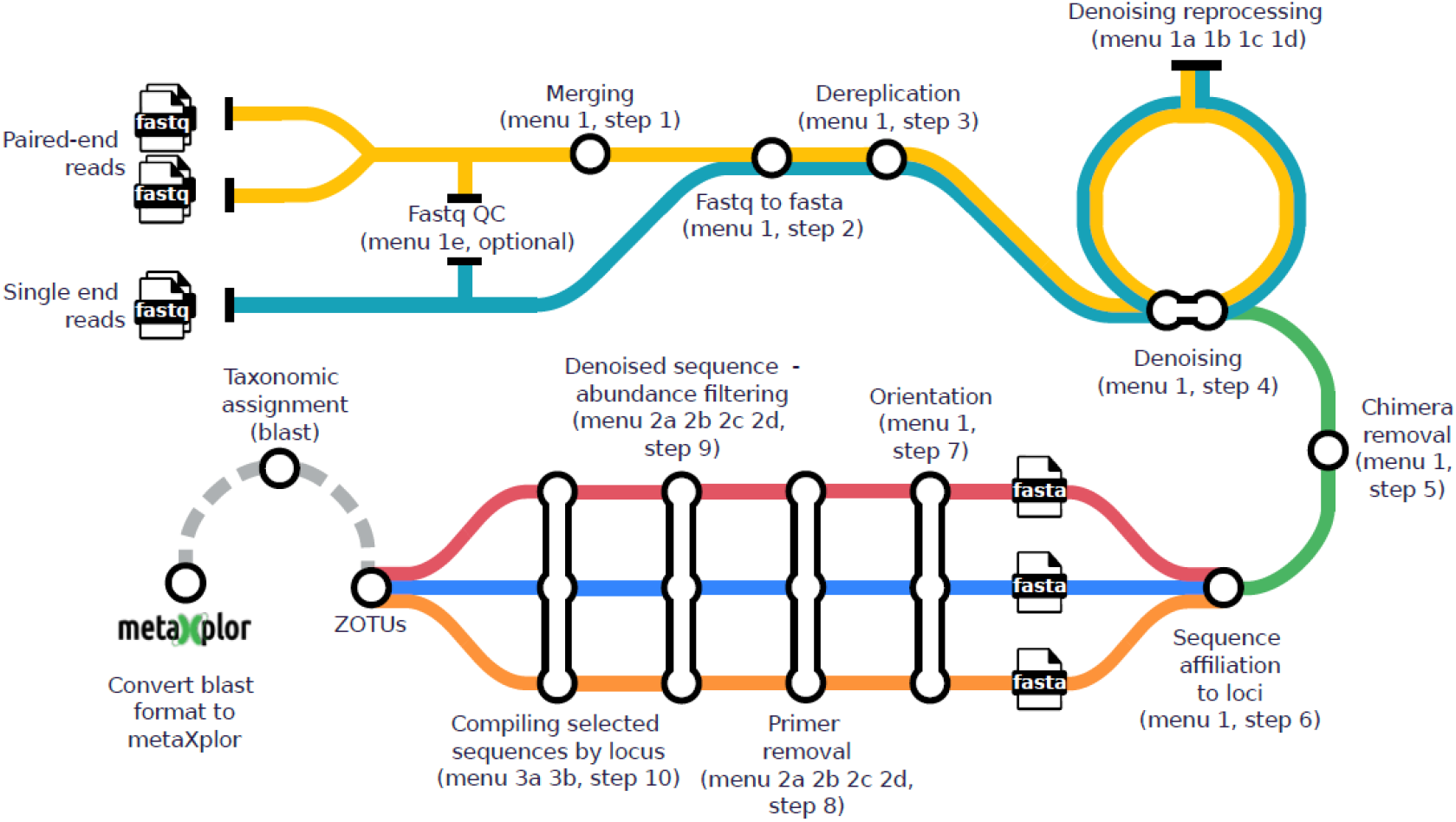
Overview of the mbctools workflow, where each station icon represents a distinct step in the process. In the initial analysis (menu 1), data processing begins with the merging for paired-end reads (step 1) and the conversion of reads from fastq to fasta (step 2). This is followed by dereplication (step 3), denoising (step 4, which can be re-run with various options via menu 1a, 1b, 1c, 1d), chimera removal (step 5), sequence affiliation to markers (step 6), and finally, sequence orientation harmonization (step 7). Following this, primer removal (step 8) and denoised sequence abundance filtering (step 9) are applied. The last step in the analysis involves obtaining per-marker fasta files containing retained ZOTUs (step 10). The menu labeled as “4” comes into play after taxonomic assignment, tuning outputs for visualization in metaXplor and ensuring long-term data accessibility.

### Prerequisites and dependencies

#### Input data types - sequencing tech & lab

*mbctools* enables the analysis of amplicon data from multiple markers, obtained from short reads (Illumina) or long reads (Oxford Nanopore Technology, PacBio). It can handle both single-end reads and paired-end reads, provided that the latter can be merged. In the eventuality where the merging of paired-end reads is prevented by either large amplicon lengths (leading to insufficient overlap zone) or the presence of low complexity sequences, *mbctools* is able to utilize only forward reads for analysis. As it relies on the underlying VSEARCH software which is highly optimized in terms of efficiency and memory utilization, it is able to accommodate large datasets deriving from numerous samples and amplicons.

#### Running environment

*mbctools* is a versatile cross-platform python package running seamlessly on Windows, Unix, or OSX systems. The installation of Python3.7 (or higher) and VSEARCH (version 2.19.0 or higher, available at https://github.com/torognes/vsearch) is required to run the pipeline, and those binaries must be added to the PATH environment variable. Additionally, for Windows users, Powershell script execution must be enabled using the *Set-ExecutionPolicy* command, as the default Windows command prompt does not provide enough flexibility to run the software. The *mbctools* package, available from the Python Package Index (PyPI) (https://pypi.org/project/mbctools/), can be installed and upgraded using the command line tool pip (*pip3 install mbctools*).

Before starting the analyses, the package requires mandatory directories and files to be pre-arranged as illustrated in Figure 3:

**Figure 3.**
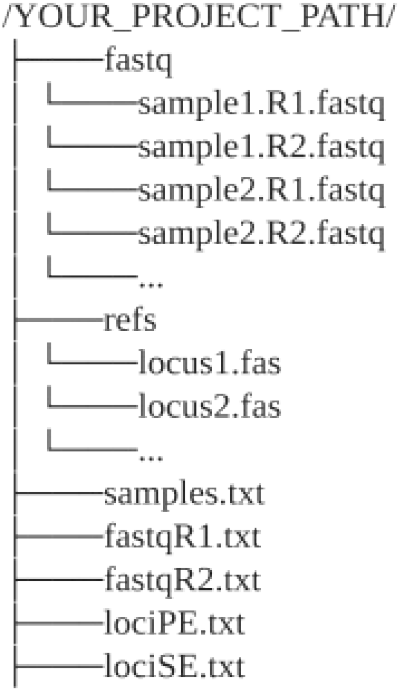
Pre-required files and directories

- A default directory, named ‘fastq’, should contain all the fastq files
- corresponding to the R1 (and R2, when working with paired-ends sequencing) reads. These files are expected to be demultiplexed, meaning that each fastq file should contain reads from a single sample.
- A default directory named ‘refs’ that contains one reference file per genetic marker in the multi-FASTA format. Each of them should contain one or more unaligned reference sequences, from the same strand and without gaps. Providing multiple reference sequences for a given marker improves diversity support and thus leads to a better assignment accuracy. In addition, the ‘sequence orientation fixing’ step runs faster when using several reference sequences. Filenames must end with the extension ‘.fas’ and must be called by the name of the corresponding marker, referred to as locus in *mbctools* (Figure 3).
- ‘lociPE.txt’ provides the list of markers (one per line) corresponding to paired-end R1/R2 reads.
- ‘lociSE.txt’ provides the list of markers (one per line) corresponding to single-end reads and for which only R1 reads will be analyzed.
- ‘samples.txt’ includes a list of sample names, where each name appears on a separate line.
- ‘fastqR1.txt’ contains the names of the *fastq* R1 files corresponding to the samples.
- ‘fastqR2.txt’ contains the names of the *fastq* R2 files corresponding to the samples.

The contents of the last three files must be sorted in the same order, i.e., line (i) of ‘samples.txt’ shall correspond to line (i) of both ‘fastqR1.txt’ and ‘fastqR2.txt’.

*mbctools* shall be launched from within the working directory in a shell console, typically by typing ‘mbctools’ if installed as a package.

### Software features and interface overview

*mbctools* features a series of menus, each containing two to four options as illustrated in Figures 4 to 8. On startup, the program displays the main menu shown in Figure 4. This menu presents descriptive keywords that enable users to easily navigate through the available menus and sections. From each menu, the user is required to select one single option at a time.

**Figure 4.**
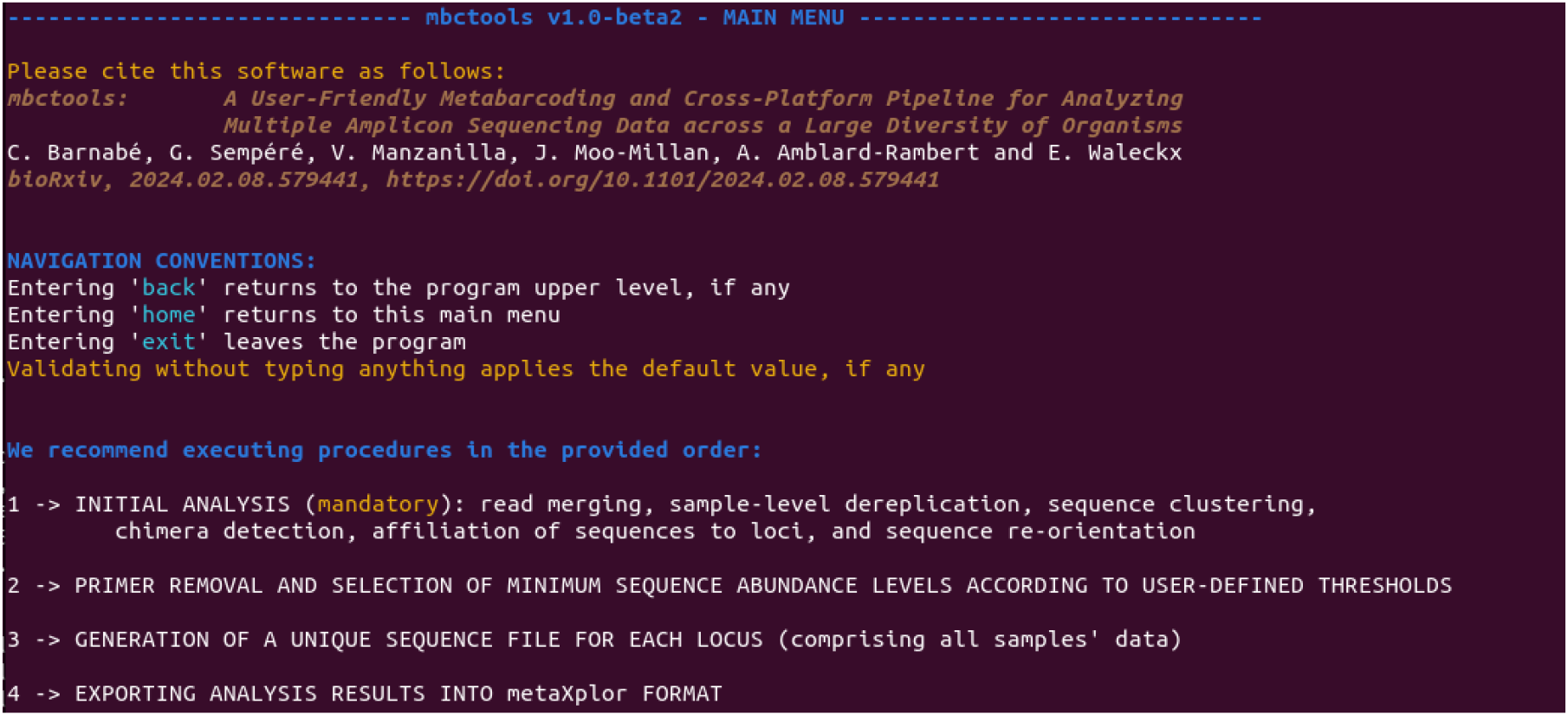
Contents of the main menu, explaining navigation conventions and providing access to the four main sections.

To analyze a new dataset, it is necessary to perform a complete initial analysis by selecting the first option from both the first and subsequent menu. The *initial analysis* processes the data through a succession of steps such as merging, sample-level dereplication, sequence denoising, chimera detection and removal, affiliation of sequences to markers (e.g., loci), and sequence re-orientation (menu 1, Figure 5). Once those are completed, primers are removed, and the denoising process may be refined by tuning the cluster size threshold (menu 2, Figure 6). Finally, centroid sequences associated with each marker can be exported to fasta files (menu 3, Figure 7). The above represents the complete cycle of data processing. Additionally (menu 4, Figure 8), *mbctools* software offers the possibility to export results in a format that is compatible with the metaXplor platform (Sempéré et al., 2021). The entire process is described as supplementary material.

**Figure 5.**
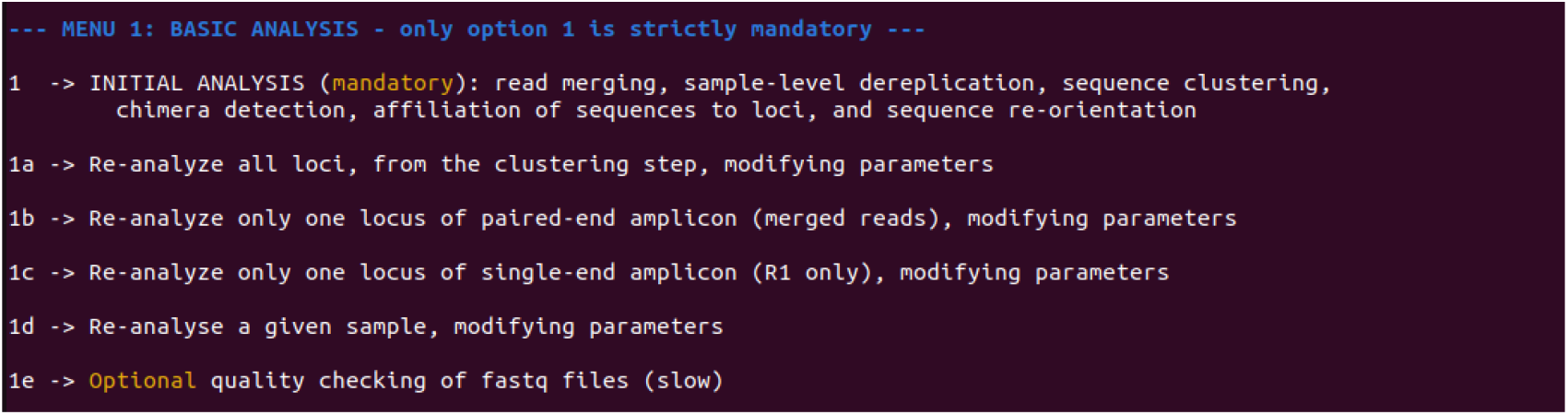
Contents of section 1 menu, in which only option 1, INITIAL ANALYSIS, is mandatory as it executes all steps required to proceed with menu 2. Options 1a to 1d may be used after option 1 in case the user wants to refine some parameters. Analysis 1e is optional and provides information about the reads’ quality.

**Figure 6.**
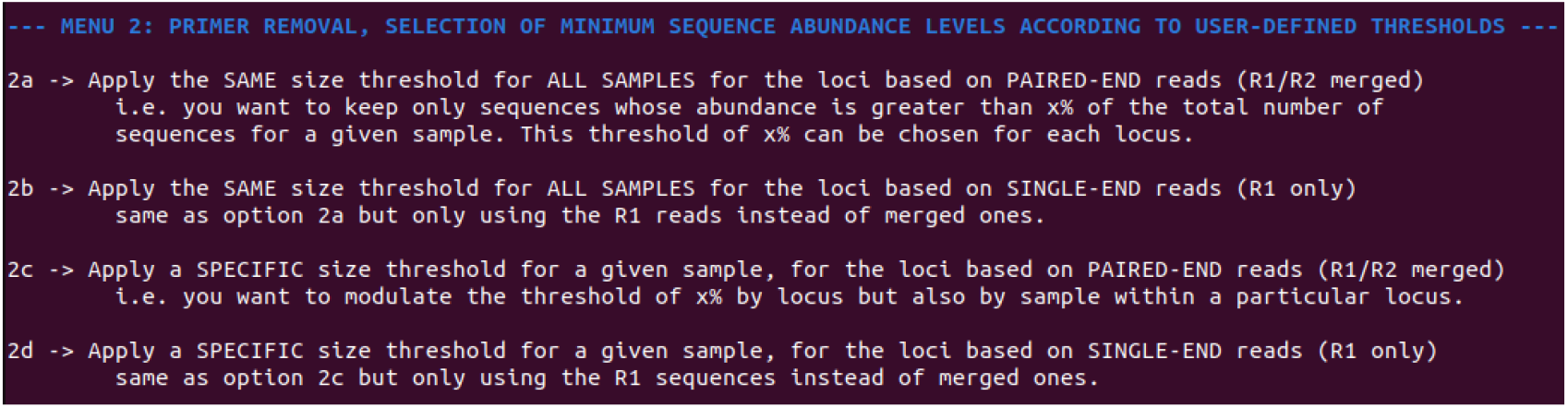
Menu 2 combines two procedures that need to be launched successively for each marker: primer removal and application of an abundance threshold for retaining centroid sequences.

**Figure 7.**
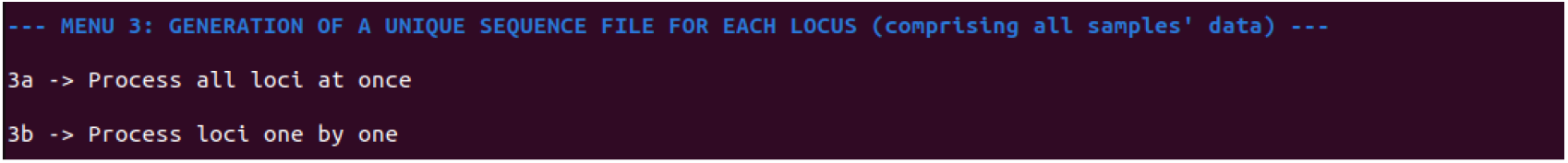
Contents of section 3 menu, allowing to obtain per-marker (per-locus) fasta files. Menu 3 provides means to generate fasta files containing the retained sequences, either for all markers (loci) at once, or for a given marker.

**Figure 8.**
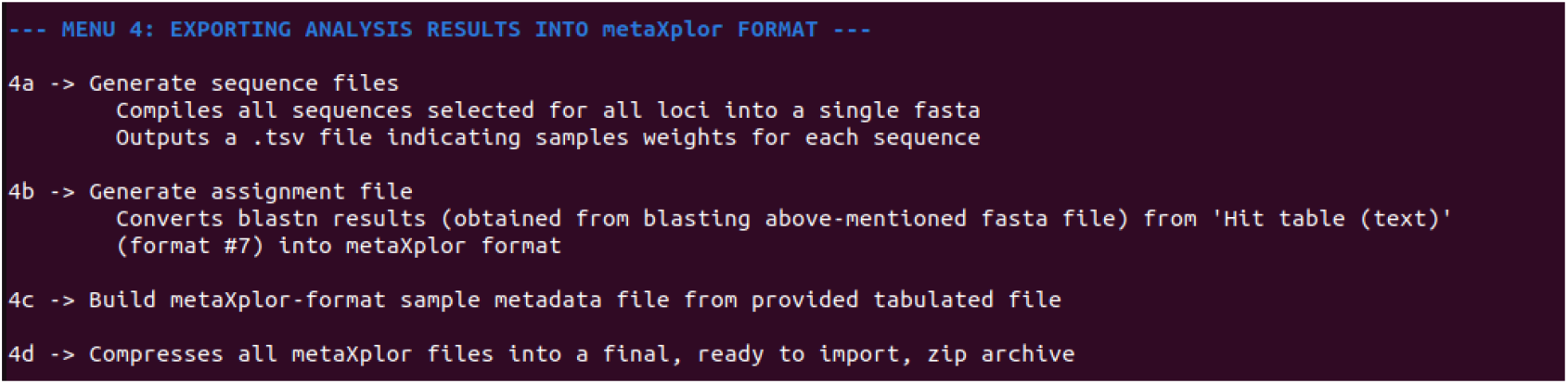
Contents of section 4 menu, guiding users in obtaining necessary files to create a metaXplor import archive (i.e., sample metadata, sequence fasta and abundance, assignment to reference taxa) in order to summarize all project results.

If, at any point, users wish to exit the program and conduct further analyses of their choice, *mbctools* allows it since it generates intermediate output files at each step.

When option 1 (initial analysis) is selected from menu 1, the menu shown on Figure 5 allows to run the initial analysis (section 1), namely read merging, sample-level dereplication, sequence clustering (denoising), chimera detection, affiliation of sequences to markers, and sequence re-orientation. In the same menu, four other sections (1a, 1b, 1c, 1d) provide means to re-process some particular markers and/or samples with adjusted parameters. In most cases the default values for those parameters will work, but there might be situations where they need to be adapted, either globally (e.g., depending on the sequencing depth, the level of similarity between targeted genetic markers), at the marker (i.e., locus) level (e.g., according to its size), or at the sample level (e.g., to account for heterogeneous sample quality). Option 1e may be used to assess the read quality for each sample, and write results to files with suffix “*.quality.txt” in the outputs directory.

When the initial analysis is completed, menu 2 should be used to remove primer sequences, and to exclude denoised sequences with an abundance considered too low for being relevant (Figure 6). This operation may be launched either in batch mode, or sample by sample in case different thresholds need to be applied.

In menu 3, the program groups sample files for each marker, creating a new file per locus with the suffix “*_allseq_select.fasta” (Figure 7). During this process, mbctools can, as an option, dereplicate sequences at the project level, ensuring that each entry in the fasta file is unique. In such cases, an additional tabulated file is generated to maintain a record of sequence abundance across the samples. Optionally generated files are then named with the following suffixes: “*_allseq_select_derep.fasta”, “*_allseq_select_derep.tsv”.

Finally, menu 4 converts *mbctools* outputs into a format suitable for importing into metaXplor (Figure 8), a web-oriented platform for storing, sharing, exploring and manipulating metagenomic pipeline outputs. metaXplor significantly contributes to enhancing the FAIR (Findable, Accessible, Interoperable, and Reusable) (Wilkinson et al., 2016) characteristics of this specific data type by serving as a robust solution for centralized data management, intuitive visualization, and long-term accessibility.

### Test data

The test data were generated in the scope of a study conducted on Chagas disease, a zoonotic parasitic illness caused by *Trypanosoma cruzi*. This disease is primarily transmitted to mammals through the feces and urine of hematophageous insects known as triatomines, commonly referred to as kissing bugs (Hemiptera: Reduviidae). *Triatoma dimidiata* is the main vector in the Yucatan Peninsula in Mexico. In 2019, the owner of an isolated rural dwelling collected thirty-three samples of *T. dimidiata* individuals from a conserved sylvatic site within the Balam Kú State Reserve, located in Campeche, Mexico (Figure 9). We used a metabarcoding approach to simultaneously characterize the genetic diversity of the vector and identify its blood meals, characterize its hindgut and midgut microbiome composition, and assess the genetic diversity of *T. cruz*i infecting *T. dimidiata* in this sylvatic ecotope. We extracted the total DNA from each *T. dimidiata* gut, and amplified different selected markers of interest for our question: (i) for *T. dimidiata* genotyping, a fragment of the vector’s Internal Transcribed Spacer 2 (ITS-2) nuclear marker was amplified with primers ITS2_200F (5′-TCG YAT CTA GGC ATT GTC TG-3′) and ITS2_200R (5-′CTC GCA GCT ACT AAG GGA ATC C-3′) previously described in (Richards et al., 2013); (ii) for the identification of vertebrate blood meals, a fragment of the vertebrate 12S rRNA gene was amplified with primers L1085 (5′-CCC AAA CTG GGA TTA GAT ACC C-3′) and H1259 (5′-GTT TGC TGA AGA TGG CGG TA-3′) (Kitano et al., 2007; Moo-Millan et al., 2019); (iii) for the identification of hindgut and midgut bacterial microbiome composition, a fragment of the bacterial 16S rRNA gene was amplified with primers E786F (5′-GAT TAG ATA CCC TGG TAG-3′) and U926R (5’-CCG TCA ATT CCT TTR AGT TT-3’) (Baker, Smith & Cowan, 2003); and (iv) to accurately genotype *T. cruzi*, we targeted markers COII, GPI, ND1 and the intergenic region of the mini-exon gene, using the primers described in (10). Then we pooled the different amplicons obtained per *T. dimidiata* specimen and sequenced them on an Illumina MiSeq. On average, we obtained 222,244 reads per sample. The reads were processed with the *mbctools* package, whose outputs were imported into metaXplor. In *mbctools*, we retained the reads with a minimum length of 100bp, clusters with a minimum size of 8 and used an alpha value of 2. The taxonomic identification of the ZOTUs was performed using BLASTn (Johnson et al., 2008). The exported CSV file with the metadata served as input for metaXplor. The analysis results are directly available on the web application under the dataset ‘mbctools_review’ (“metaXplor: mbctools data set,” 2024) (Figure 9).

**Figure 9.**
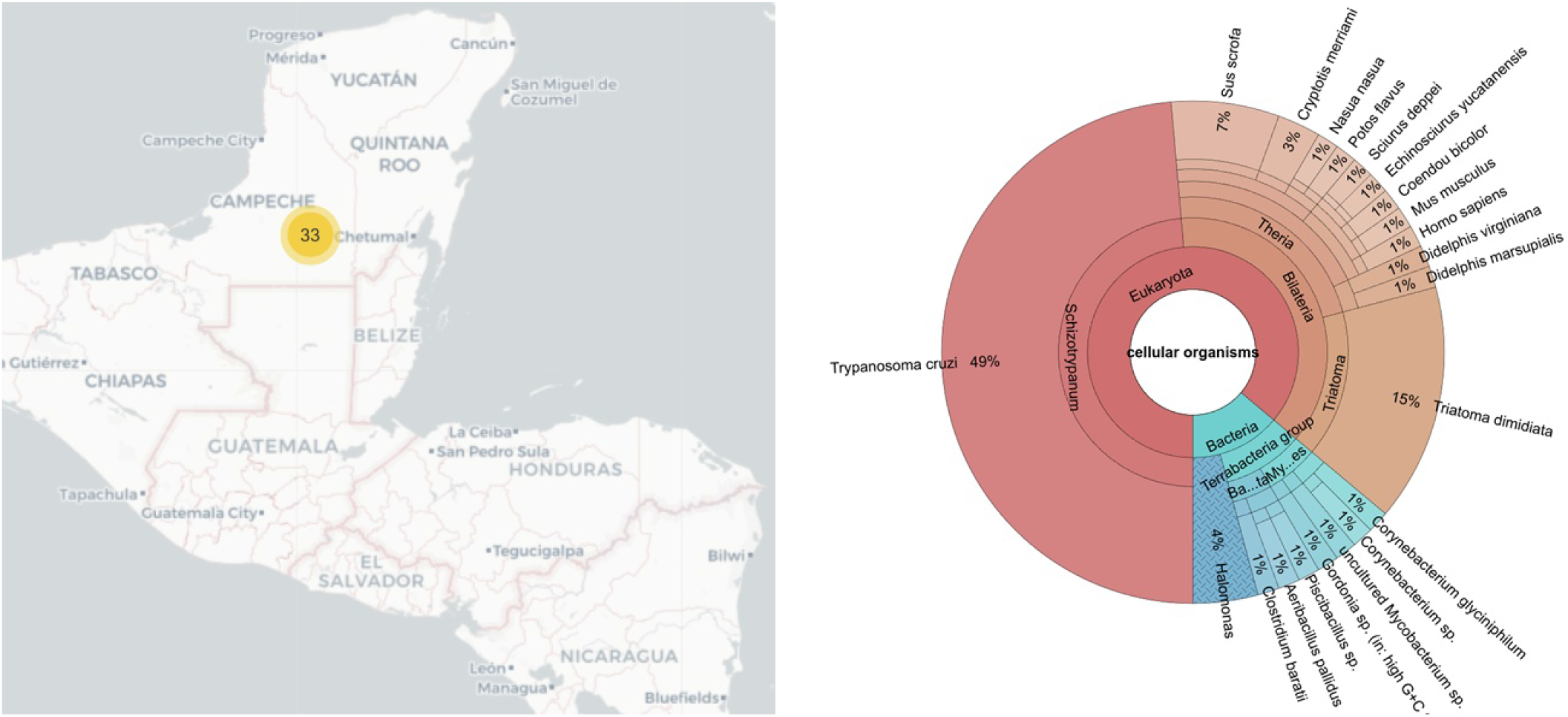
Graphical output from the metaXplor web interface. On the left side the geographic origin of the thirty three samples. On the right side the taxonomic proportion of each identified organism

We identified an average of 33 ZOTUs per *T. dimidiata* sample, and only 9.61% of the reads were discarded during the analysis (Table 1). After analysis, we found thirteen blood-meal sources (the most frequently identified being *Homo sapiens, Sus scrofa* and *Mus musculus*), six bacterial Orders (Bacillales, Oceanospirillales and Micrococcales being the most abundant), along with *T. cruzi* and *T. dimidiata* (Figure 9). The test data input and output files are freely accessible on the DataSuds repository (Manzanilla et al., 2024).

**Table 1.** Statistics concerning the processing of the test data set with the number of reads per sample and the attribution to each marker.

| Sample | Reads per sample | Merging | Number of clusters | clusters with no chimera | Cluster per marker |  |  |  |  |  |  |  |
| --- | --- | --- | --- | --- | --- | --- | --- | --- | --- | --- | --- | --- |
|  |  |  |  |  | 12S | 16S | COII | GPI | ITS2 | ND1 | MINEX | MINEX_SE |
| CON001 | 215137 | 0.8696 | 11 | 11 | 0 | 15 | 0 | 0 | 2 | 3 | 6 | 0 |
| CON002 | 268415 | 0.9229 | 17 | 13 | 18 | 12 | 0 | 0 | 2 | 0 | 0 | 0 |
| CON003 | 242823 | 0.8636 | 87 | 87 | 0 | 9 | 0 | 1 | 2 | 0 | 12 | 0 |
| CON004 | 205358 | 0.9344 | 3 | 3 | 0 | 11 | 0 | 0 | 3 | 0 | 0 | 0 |
| CON005 | 149945 | 0.9408 | 7 | 7 | 1 | 6 | 0 | 0 | 2 | 0 | 0 | 0 |
| CON006 | 180763 | 0.9311 | 2 | 2 | 0 | 11 | 0 | 0 | 3 | 0 | 0 | 0 |
| CON007 | 7898 | 0.891 | 2 | 2 | 1 | 0 | 0 | 0 | 2 | 0 | 0 | 0 |
| CON008 | 200589 | 0.924 | 3 | 3 | 0 | 19 | 0 | 0 | 2 | 0 | 0 | 0 |
| CON009 | 261914 | 0.9226 | 2 | 2 | 0 | 15 | 0 | 0 | 2 | 0 | 0 | 0 |
| CON010 | 283721 | 0.9325 | 139 | 126 | 16 | 16 | 0 | 0 | 2 | 0 | 0 | 0 |
| CON011 | 186936 | 0.9112 | 1 | 1 | 0 | 16 | 0 | 0 | 2 | 0 | 0 | 0 |
| CON012 | 194776 | 0.8326 | 61 | 60 | 14 | 10 | 2 | 1 | 2 | 0 | 24 | 4 |
| CON013 | 189441 | 0.9303 | 14 | 14 | 4 | 8 | 0 | 0 | 4 | 0 | 5 | 0 |
| CON014 | 151880 | 0.9086 | 25 | 25 | 0 | 8 | 0 | 0 | 30 | 0 | 0 | 0 |
| CON015 | 221302 | 0.9174 | 1 | 1 | 0 | 14 | 0 | 0 | 2 | 0 | 0 | 0 |
| CON016 | 139546 | 0.8699 | 46 | 46 | 2 | 6 | 2 | 1 | 10 | 0 | 14 | 3 |
| CON017 | 228038 | 0.8812 | 3 | 3 | 0 | 11 | 0 | 0 | 2 | 0 | 0 | 0 |
| CON018 | 258481 | 0.9012 | 28 | 27 | 6 | 12 | 0 | 0 | 15 | 0 | 0 | 0 |
| CON019 | 251403 | 0.7904 | 77 | 77 | 1 | 14 | 4 | 2 | 10 | 0 | 3 | 0 |
| CON020 | 223972 | 0.9283 | 24 | 24 | 2 | 1 | 0 | 0 | 4 | 0 | 0 | 0 |
| CON021 | 309618 | 0.9346 | 8 | 8 | 0 | 15 | 0 | 0 | 14 | 0 | 0 | 0 |
| CON022 | 197125 | 0.8479 | 83 | 83 | 11 | 8 | 2 | 2 | 4 | 0 | 5 | 0 |
| CON023 | 193857 | 0.7558 | 53 | 53 | 1 | 7 | 2 | 1 | 5 | 5 | 15 | 0 |
| CON024 | 190310 | 0.9132 | 62 | 62 | 13 | 20 | 0 | 0 | 2 | 0 | 0 | 0 |
| CON025 | 273603 | 0.9104 | 44 | 37 | 29 | 15 | 0 | 0 | 2 | 0 | 0 | 0 |
| CON026 | 284204 | 0.945 | 2 | 2 | 1 | 17 | 0 | 0 | 2 | 0 | 0 | 0 |
| CON027 | 436822 | 0.9225 | 5 | 5 | 1 | 34 | 0 | 0 | 2 | 0 | 0 | 0 |
| CON028 | 243235 | 0.9385 | 63 | 63 | 1 | 12 | 0 | 0 | 4 | 0 | 0 | 0 |
| CON029 | 219907 | 0.9486 | 13 | 13 | 4 | 11 | 0 | 0 | 2 | 0 | 0 | 0 |
| CON030 | 306738 | 0.9456 | 17 | 17 | 0 | 10 | 0 | 0 | 26 | 0 | 0 | 0 |
| CON031 | 172815 | 0.9398 | 87 | 87 | 1 | 11 | 0 | 0 | 2 | 0 | 0 | 0 |
| CON032 | 206078 | 0.9069 | 34 | 33 | 1 | 13 | 0 | 0 | 24 | 0 | 0 | 0 |
| CON033 | 237429 | 0.9158 | 92 | 92 | 37 | 14 | 0 | 0 | 2 | 0 | 0 | 0 |
| Average | 222244.8 | 0.9 | 33.8 | 33 | 5 | 12.2 | 0.36 | 0.24 | 5.88 | 0.24 | 2.55 | 0.21 |

## Conclusion

*mbctools* facilitates the processing of amplicon datasets for metabarcoding studies by offering menus that guide users through the various steps of the data analysis. It is, to our knowledge, the only tool eliminating the need for the computer literacy normally required to utilize command-line based software as complex as the underlying VSEARCH, while allowing to process multiple genetic markers simultaneously. *mbctools* offers the possibility to reprocess specific markers and samples with customized parameters, providing means to adapt to the diversity of metabarcoding datasets. In the end, it generates *fasta* files composed of the filtered ZOTUs ready for the taxonomic assignment step. Although BLASTn is a commonly used and straightforward solution for this task, given the diversity of tools and approaches (k-mer based Kraken2 (Wood, Lu & Langmead, 2019), Kaiju (Menzel, Ng & Krogh, 2016), phylogeny-based PhyloSift (Darling et al., 2014)), we leave to the user to proceed with taxonomic assignment, based on his own practice or the literature (18). To promote data management, visualization and long-term accessibility, *mbctools* offers a file conversion functionality compatible with the metaXplor web application which aims at centralizing online meta-omic data, while offering user-friendly means to interact with it. The metaXplor instance hosted at CIRAD (https://metaxplor.cirad.fr/) is indeed used by several teams to keep track of previous project data (be it private or public) and proves useful in helping scientists quickly recover precise information. As an example, metaXplor was utilized in the scope of 24 shotgun metagenomics projects based on VANA (Virion-Associated Nucleic Acids), conducted to uncover viral diversity (Moubset et al., 2022). It allowed for the incremental building of a data repository that efficiently manages and provides means to analyze the extensive sequence datasets thus generated. This platform facilitated similarity-based searches and phylogenetic analyses, significantly enhancing the retrieval and re-analysis of data, thereby promoting viral discovery and classification. Although we initially developed the *mbctools* pipeline and software for reproducibility purposes across our teams working on infectious diseases, the pipeline’s strength resides in its capacity to handle diverse data and be tailored to different research needs. This tool plays a central role in training sessions provided by our teams in developing countries, and its functionalities and user-friendliness are recurrently extended according to the feedback they generate. Depending on the success of this first release, future versions may add support for new sequencing technologies, embedded taxonomic assignment solutions or phylogenetic tree building.

## Authors’ contributions

C.B and G.S designed and conceived the software based on E.W.’s specifications. V.M. did the Python packaging. E.W, G.S, V.M, A.A-R and C.B wrote the article. A.A-R. provided the data to test mbctools. All authors read and approved the final manuscript.

## Data, scripts, code, and supplementary information availability

*mbctools* is openly accessible on Github at https://github.com/GuilhemSempere/mbctools and on PyPI at https://pypi.org/project/mbctools/. The main pipeline description is available as supplementary material, as are higher-definition versions of Figures 1 to 3.

## Acknowledgements

The authors would like to thank Frédéric Mahé for taking the time to introduce VSEARCH functionalities, as well as Dorian Grasset for testing the OSX version and creating a GitHub action to automatize PyPI package generation. Preprint version 2 of this article has been peer-reviewed and recommended by Peer Community In Genomics (https://doi.org/10.24072/pci.genomics.100370; Pollet, 2024).

