## Supplementary material for "mbctools: A User-Friendly Metabarcoding and Cross-Platform Pipeline for Analyzing Multiple Amplicon Sequencing Data across a Large Diversity of Organisms": Main pipeline description (from another version of this preprint: version 3, 2024-09-21)

†Co–first authors

### Annex 1: Main pipeline description

#### *From fastq files to relevant amplicon sequences (menu 1)*

The filenames in the examples given in this section follow a number of conventions:

- *occurrences of the string "(paired|single)" actually represent either "paired" or "single" depending on the selection made by the user*
- *occurrences of the character string "(L1|L2)" actually represent the name of the locus selected by the user*
- *occurrences of the character string "(Sample1|Sample2)" actually represent the name of the sample selected by the user*
- *dir_fastq: directory containing the fastq files*
- *fastqR1: name of the fastq file containing the R1 reads*
- *fastqR2: name of the fastq file containing the R2 reads*

The workflow that applies to the sequencing data is detailed below in seven steps:

##### **Step 1: conversion of R1 / single-end fastq file into a fasta file.**

Using VSEARCH, this command converts the single-end fastq file into a multi-fasta file.

Underlying VSEARCH command:

*vsearch --fastq_filter dir_fastq/fastaqR1 –fastaout ../tmp_files/(Sample1|Sample2)_singleEnd.fa*

##### **Step 2: merging**

When the length of the expected amplicon allows a sufficient overlap, this step merges R1/R2 paired-end *fastq* files into a *fasta* file. When the merged reads are lower than 300bp, it removes the read tails (--*fastq_allowmergestagger*).

Underlying VSEARCH command:

*vsearch --fastq_mergepairs dir_fastq/fastqR1 --reverse dir_fastq/fastqR2 --fastaout
 ./tmp_files/(Sample1|Sample2)_pairedEnd.fa --fastq_allowmergestagger --relabel sample=(Sample1|Sample2)_merged.*

##### **Step 3: dereplication**

This step aggregates identical sequences into a unique sequence from both strands for R1/R2 paired-end merged sequences and single-end R1 sequences (if it applies) into a *fasta* file where we keep track of sequence abundance in the header.

Underlying VSEARCH command:

*vsearch --fastx_uniques ../tmp_files/(Sample1|Sample2)_(paired|single)End.fa  --fastaout ../tmp_files/(Sample1|Sample2)_(paired|single)End_derep.fas --sizeout --strand both*

##### **Step 4: Denoising**

Denoising of amplicons following the UNOISE version3 algorithm by Robert Edgar (Edgar, 2016b), and clustering similar sequences around a centroid (i.e. the sequence with higher abundance in its cluster).

Underlying VSEARCH command:

*vsearch --cluster_unoise ../tmp_files/(Sample1|Sample2)_(paired|single)End_derep.fas --sizein --centroids*

*../tmp_files/(Sample1|Sample2)_(paired|single)End_cluster.fas --strand both --minsize int --sizeout --unoise_alph int  --minseqlength int*

##### **Step 5: chimera detection and removal**

Detection and removal of potential chimeras in denoised sequences, using the UCHIME2 algorithm (Edgar, 2016a).

Underlying VSEARCH command:

*vsearch --uchime3_dsenovo ../tmp_files/(Sample1|Sample2)_(paired|single)End_cluster.fas --nonchimeras ../tmp_files/(Sample1|Sample2)_(paired|single)End_cluster_OK.fas*

##### **Step 6: affiliation**

To assign each denoised sequence to a specific investigated marker, a comparison is conducted between reference sequences of the  targeted markers (specified by the "--db" argument) and the cluster-centroid sequences from each sample. This comparison is performed using a global pairwise alignment method, allowing to determine the affiliation of each denoised sequence to its respective marker.

Underlying VSEARCH command:

*vsearch --usearch_global ../tmp_files/(Sample1|Sample2)_(paired|single)End_cluster_OK.fas --db ../refs/(L1|L2).fas*

*--matched  ../loci/(L1|L2)/(Sample1|Sample2)_(paired|single)End.fas --id 70 --strand both*

##### **Step 7: orientation**

When necessary, the orientation of remaining sequences is corrected to be the same as the reference sequences.

Underlying VSEARCH command:

*vsearch --orient ../loci/(L1|L2)/(Sample1|Sample2)_(paired|single)End.fas --db ../refs/(L1|L2).fas –fastaout ../loci/(L1|L2)/(Sample1|Sample2)_(paired|single)End_orient.fas*

The essential inputs for options under menu 1 involve directing the system to all the required files and supplying a set of parameters (some of which having default values), to customize the processing steps performed by VSEARCH. These inputs are prompted from the shell environment.

- Full path of the folder containing *fastq* files (default, *./fastq*) where “.” refers to the current directory
- Name of the file listing R1 *fastq* (plain, or gzipped if installed VSEARCH version supports it) filenames (default, *fastqR1.txt*). *NB: file lists are all expected to provide one element per line*
- Name of the file listing R2 *fastq* (plain, or gzipped if installed VSEARCH version supports it) filenames (default, *fastqR2.txt*).
   *NB: Target file must exist but may be empty to process only single-end or unmerged R1 reads*
- Name of the file listing the names of the loci expected to be found in paired-end reads (default, *lociPE.txt*).
   *NB: Target file must exist but may be empty to process only single-end or unmerged R1 reads*
- Name of the file listing the names of the loci expected to be found in single-end reads (default, *lociSE.txt*)
   *NB: Target file must exist but may be empty to process only merged paired-end reads*
- Name of the file listing sample names (default, *samples.txt*)
- Minimum abundance (VSEARCH parameter named *--minsize)* for retaining denoised sequences (default, *8*)
- Minimum length (VSEARCH parameter named *--minseqlength*) of retained sequences for any locus (default, *100*)
- Value for the alpha parameter (VSEARCH parameter named *--unoise_alpha*) used in the UNOISE [REF] clustering algorithm (default, *2*)
- Identity percentage parameter (VSEARCH parameter named *--id*) to match the denoised sequences against local references (default, *70%*)

Note: Options 1a to 1d allow refining the 4 last parameters (the ones used for tuning VSEARCH behavior) in order to apply particular settings on all or some of the samples and loci. Previously dereplicated sequences are then used again for re-processing selected data from the denoising/clustering step #4.

All the specified parameters and their values are automatically stored in the following configuration file: *outputs/parameters_option_1.cfg*. Once this file has been created, it can be stored separately and reused to initiate the main analysis directly, eliminating the need to re-enter each parameter individually. This can be accomplished as follows by passing its full path as an argument to mbctools, as follows: *mbctools outputs/parameters_option_1.cfg.* Using the software in this manner transforms it into a purely command-line tool, eliminating the necessity for direct interaction with the interface and facilitating its integration into HPC pipelines.

#### *Primer removal and abundance filtering (menu 2)*

This section aims at cleaning up amplicon sequences by removing primers from their ends, and ignoring sequences whose abundance is considered too rare to be relevant. The following user inputs are requested for options under menu 2:

- Locus to work with (no default, to choose among those featured in the project)
- Length of both right and left primers to trim from the final selected centroid sequences (no default)
- Minimum abundance threshold to use for the selected locus, e.g. entering 5 will ignore denoised sequences covering under 5% of the sample’s total reads
- The name of the sample to work with (only applies to options 2c and 2d since previous ones process all samples)

#### *Compiling selected sequences by locus (menu 3)*

Once option 2 has been executed, users may want to gather all samples’ relevant ZOTUsinto one file per locus, typically for further analyses such as phylogeny.

The only expected user input is then the locus for which to group all samples’ selected sequences into a single multi-fasta file called *locusX_allseq_select.fasta*.

#### *Execution outputs*

During the execution of basic analyses, several directories are automatically generated. These include the "loci" directory, which consists of subdirectories named after the corresponding reference files for each locus. Another directory named "scripts" is created to store intermediate VSEARCH invocation scripts specific to the current project, generated based on the selected parameters. Additionally, a directory named "tmp_files" is generated to hold all intermediate temporary files produced during various procedures. Lastly, an "outputs" directory is created, containing quality checking statistics, execution logs, and settings files related to the tasks initiated from the different menus.

For each sample, up to six output files  generated in the corresponding subfolders of the *loci* directory:

- *(Sample1|Sample2)_(paired|single)End.fas* corresponding to outputs of options from menu 1,
- *(Sample1|Sample2)_(paired|single)End_orient.fas*, equivalent of the latter but with all sequences oriented in the same strand,
- *(Sample1|Sample2)_paired(single)End_select.fas*, corresponding to sequences retained after applying a particular abundance threshold (options from menu 2).

When running concatenation option (#3) the above-mentioned additional file called *locusX_allseq_select.fasta* is generated for each locus, grouping all finally retained sequences for that locus, and which can be used directly in phylogenetic programs.

Exporting analysis results into metaXplor format

Menu 4 guides users into generating a zip archive ready to import into metaXplor:

- option 4a dereplicates centroid sequences at the project level, keeping track of sample abundance via a .tsv file
- option 4b, providing that BLASTn has been run appropriately on the above, converts its results into a metaXplor assignment file
- option 4c takes a tabulated sample metadata file and turns it into the appropriate format
- option 4d compresses files generated from previous steps into a single zip archive, ready for import into metaXplor

While processing data, the system keeps track of VSEARCH execution statistics, logging information and configuration parameters in the *outputs* folder, in files named after the executed procedure  (e.g. for the main analysis #1, *Stats_option_1.txt,* , *res1.log* and *parameters_option_1.cfg* respectively).

### Annex 2: VSEARCH parameters mentioned in Annex 1

--centroids ‘filename’: Output cluster centroid sequences to filename, in fasta format. The centroid is the sequence that seeded the cluster (i.e. the first sequence of the cluster).

--db ‘filename’: indicates the reference file to use for global pairwise alignment.

--fastq_allowmergestagger: this option allows to merge staggered read pairs. Staggered pairs are pairs where the 3’ end of the reverse read has an overhang to the left of the 5’ end of the forward read. This situation can occur when a very short fragment is sequenced. The 3’ overhang of the reverse read is not included in the merged sequence.

--id real: sets the minimum pairwise sequence identity to use in the command; value in range [0 – 1.0].

--matched ‘filename’: writes sequences matching database target to filename, in fasta format.

--minseqlength int: retains length sequences >= integer.

--minsize int: retains abundances >= integer; default is 8.

--nonchimeras ‘filename’: outputs the fasta file ‘filename’ containing all non-chimeric sequences.

--orient ‘filename’: orients the sequences in the given file in the same sense as in the reference file specified using --db.

--sizein: takes into account abundance annotations of the headers of input file.

--sizeout: adds abundance annotations to output fasta file headers.

--strand both: checks both strands when comparing sequences.

--uchime3_denovo: improved detection of chimeric sequences.

--unoise_alph int: sets the *alpha* parameter used in the *--cluster_unoise* command; default is 2.0
